# One leaf for all: Chemical traits of single leaves measured at the leaf surface using Near infrared-reflectance spectroscopy (NIRS)

**DOI:** 10.1101/2020.03.26.010462

**Authors:** Matteo Petit Bon, Hanna Böhner, Sissel Kaino, Torunn Moe, Kari Anne Bråthen

## Abstract

1. The leaf is an essential unit for measures of plant ecological traits. Yet, measures of plant chemical traits are often achieved by merging several leaves, masking potential foliar variation within and among plant individuals. This is also the case with cost-effective measures derived using Near-infrared reflectance spectroscopy (NIRS). The calibration models developed for converting NIRS spectral information to chemical traits are typically based on spectra from merged and milled leaves. In this study we ask if such calibration models can be applied to spectra derived from whole leaves, providing measures of chemical traits of single leaves.
2. We sampled cohorts of single leaves from different biogeographic regions, growth forms, species and phenological stages in order to include variation in leaf and chemical traits. For each cohort we first sampled NIRS-spectra from each whole, single leaf, including leaf sizes down to Ø 4 mm (the minimum area of our NIRS application). Next, we merged, milled and tableted the leaves and sampled spectra from the cohort as a tablet. We applied arctic-alpine calibration models to all spectra and derived chemical traits. Finally, we evaluated the performance of the models in predicting chemical traits of whole, single leaves by comparing the traits derived at the level of leaves to that of the tablets.
3. We found that the arctic-alpine calibration models can successfully be applied to single, whole leaves for measures of Nitrogen (R^2^=0.88, RMSE=0.824), Phosphorus (R^2^=0.65, RMSE=0.081), and Carbon (R^2^=0.78, RMSE=2.199) content. For Silicon content we found the method acceptable when applied to Silicon-rich growth forms (R^2^=0.67, RMSE=0.677). We found a considerable variation in chemical trait values among leaves within the cohorts.
4. This time- and cost-efficient NIRS-application provides non-destructive measures of a set of chemical traits in single, whole leaves, including leaves of small sizes. The application can facilitate research into the scales of variability of chemical traits and include intraindividual variation. Potential trade-offs among chemical traits and other traits within the leaf unit can be identified and be related to ecological processes. In sum this NIRS-application can facilitate further ecological understanding of the role of leaf chemical traits.

## Introduction

The essential role of chemical constituents in plants and ecosystem functioning is repeatedly emphasized (White 1993; Elser *et al.* 1996; Aerts & Chapin 2000; Güsewell 2004; Elser *et al.* 2007; LeBauer & Treseder 2008; Cebrian *et al.* 2009; Elser *et al.* 2010; Fay *et al.* 2015). Foliar chemical constituents show interspecific variability at both spatial and temporal scales within and across ecosystems, and are closely related to plant performance and ecological interactions (Güsewell 2004). Furthermore, foliar chemical content is among the plant traits with the highest intraspecific variability (Albert *et al.* 2010; Siefert *et al.* 2015; Fajardo & Siefert 2016), that may also include intraindividual variability (Ely *et al.* 2019). For instance, trait measures at the leaf level have been found to explain a considerable part of trait variability among tropical trees (Messier, McGill & Lechowicz 2010), large leaved food plants (Ely *et al.* 2019) and alpine plants (Albert *et al.* 2010). Albert *et al.* (2010) found intraindividual trait variation to be the largest for leaf dry matter content (LDMC) (ratio of dry to fresh leaf mass) in the dwarf shrub *Vaccinium myrtillus*. LDMC variability of 8 %, 37 % and 55 % was explained by differences among populations, individuals and leaves respectively. However, most methods to measure foliar chemical content require more than a single leaf, especially when working with small arctic and alpine plant species, causing knowledge about intraspecific variability in chemical traits to be at a rudimentary stage. Yet, the single leaf is a plant unit involved in ecological interactions. Thus, chemical trait information at the leaf level is likely to prove useful both to eco-physiological understanding of trade-offs within plants, and to how chemical traits and their variability are likely to affect ecosystem process rates (Bolnick *et al.* 2003; Albert *et al.* 2011; Bolnick *et al.* 2011). In this study, we address time- and cost-efficient methodology for measuring chemical traits in single leaves, a scale of potential relevance to several ecological questions.

Near infra-red reflectance spectroscopy (NIRS) has recently been found to provide cost-efficient and accurate measures of leaf chemical traits independent of species, phenology, ecological context and region (Serbin *et al.*; Petisco *et al.* 2005; Serbin *et al.* 2014; Smis *et al.* 2014; Couture *et al.* 2016; Murguzur *et al.* 2019). Such cost-efficient measures open avenues for incorporating leaf chemical traits in large-scale ecological studies. This is strengthened by the fact that a single measure, one spectrum, of a sample is enough for predicting several chemical traits. Furthermore, the sampling of a spectrum is non-destructive, causing analyzed plant material to be available for further studies of, for example, the content of other constituents or of ecological processes such as decomposition rates. However, typical NIRS-applications are based on calibration models between NIRS derived spectra of dried and milled leaves versus their analytically derived chemical content. In order to have enough milled leaf material it is often necessary to merge several leaves, especially of species with small leaves, masking potential chemical trait variability among leaves. And although cost-efficient, the process-time for milling is still a constraint (Couture *et al.* 2016). Here we ask if arctic-alpine calibration models for NIRS-based measures of chemical traits (Smis *et al.* 2014; Murguzur *et al.* 2019), models that are based on NIRS-spectra from dried, milled and tableted leaves, can be applied to NIRS-spectra of single, whole leaves and still provide accurate measures of chemical content.

The precision of NIRS calibration models for measures of chemical constituents is dependent on the precision and bias of the analytical techniques from which the chemical constituents are retrieved and the NIRS-spectra are fitted (Chodak 2008). NIRS calibration models can thus only be as precise as the chemical analysis methods upon which they are based (Coates 2002). Any analytical technique imprecision, although within the acceptable range of precision requirements that apply to standard method performance for analytical methods (AOAC International 2016), can reduce the fit between the actual constituent values and the NIRS-spectra (Coates 2002). Because precision requirements are lower for small contents (Horwitz & Albert 2006), the fit can be poorer for nutrients with small content. Furthermore, any bias, i.e. a systematic shift in measured quantity above or below the true content, will reduce the fit with spectra derived from NIRS. It is therefore a great challenge to assess the actual accuracy of NIRS based measures. Still, NIRS calibration models for a range of chemical constituents perform well, and are not inferior to chemical analysis in terms of accuracy (Smis *et al.* 2014; Murguzur *et al.* 2019). The arctic-alpine NIRS calibration models are developed for measures of foliar Nitrogen, Phosphorus, Carbon (Murguzur *et al.* 2019), and Silicon content (Smis *et al.* 2014). We chose these models for this particular study because they provide accurate measures of chemical traits for a range of growth forms and species, at a range of phenological stages in both arctic and alpine environments. Hence, they can potentially provide robust measures of chemical content of single leaves from any species in these environments. Furthermore, there is potential to build on these arctic-alpine NIRS calibration models to become global (Murguzur *et al.* 2019), making them relevant to a range of other environments.

For our testing, we sampled cohorts of leaves from a range of plant individuals from three biogeographic regions, from different vegetation types, growth forms, species and phenological stages. With this wide range of leaf-cohorts we aimed to maximize the range of leaf types, and the range of foliar chemical content of our samples, according to guidelines for how to develop optimally performing methods (AOAC International 2016). Within each leaf-cohort, we dried the leaves, sampled spectra and predicted the chemical content of each single, whole leaf. We then calculated the average content per cohort and compared this average to the prediction achieved from the same cohort in the form of a tablet (all the leaves of the cohort merged, milled and pressed into a tablet). We also assessed to what extent the single leaves within a cohort showed variation in their chemical traits, including stoichiometric relations. We hypothesized that the arctic-alpine NIRS calibration models performed well for the prediction of chemical trait values of single, whole leaves. We also hypothesized that the chemical trait values differed among the single leaves within the leaf-cohorts.

## Methods

### Leaf sampling

The sampling was conducted on Svalbard, in Finnmark and in Troms (Norway), representing the biogeographic regions of the high-Arctic, the sub-Arctic alpine and the Boreal-alpine respectively. The sampling in Svalbard was conducted in Adventdalen (78° 10′ N, 16° 05′ E), a wide, formerly glaciated valley on the island of Spitsbergen, during the summer of 2016. We sampled leaves in dry heaths, mesic heaths, and wetlands, which represent the majority of habitat-types found across the archipelago (Elvebakk 2005). Both Finnmark and Troms belong to the Norwegian part of Fennoscandia. The sampling in Finnmark was conducted in the low alpine zone at 300–400 m a.s.l. at Ifjordfjellet (70° 27′ N, 27° 08′ E), during the summer season of 2015. The region is mainly characterized by dwarf-shrub heaths (Walker *et al.* 2005), whereas we sampled leaves mainly from tundra grasslands that typically dominate river plains and that host a wide variety of growth forms. The sampling in Troms was conducted in the low alpine zone at 400-500 m a.s.l. in the mountainous areas surrounding the city of Tromsø (69° 40′ N, 18° 55′ E) during the summer of 2017. Additional sampling of senescent leaves and litter, hereafter denoted as leaf litter, was conducted in the fall in 2017 in the boreal forest of Troms at approx. 50-100 m a.s.l.. We collected a total of 1677 fresh leaves for a total of 97 leaf-cohorts (set of single leaves merged into tablets), and we collected leaf litter for a total of 20 litter-cohorts (without separating between single leaves) (Table 1). Within each biogeographic region the cohorts were collected from different vegetation types, growth forms, species and from different phenological stages.

**Table 1.**
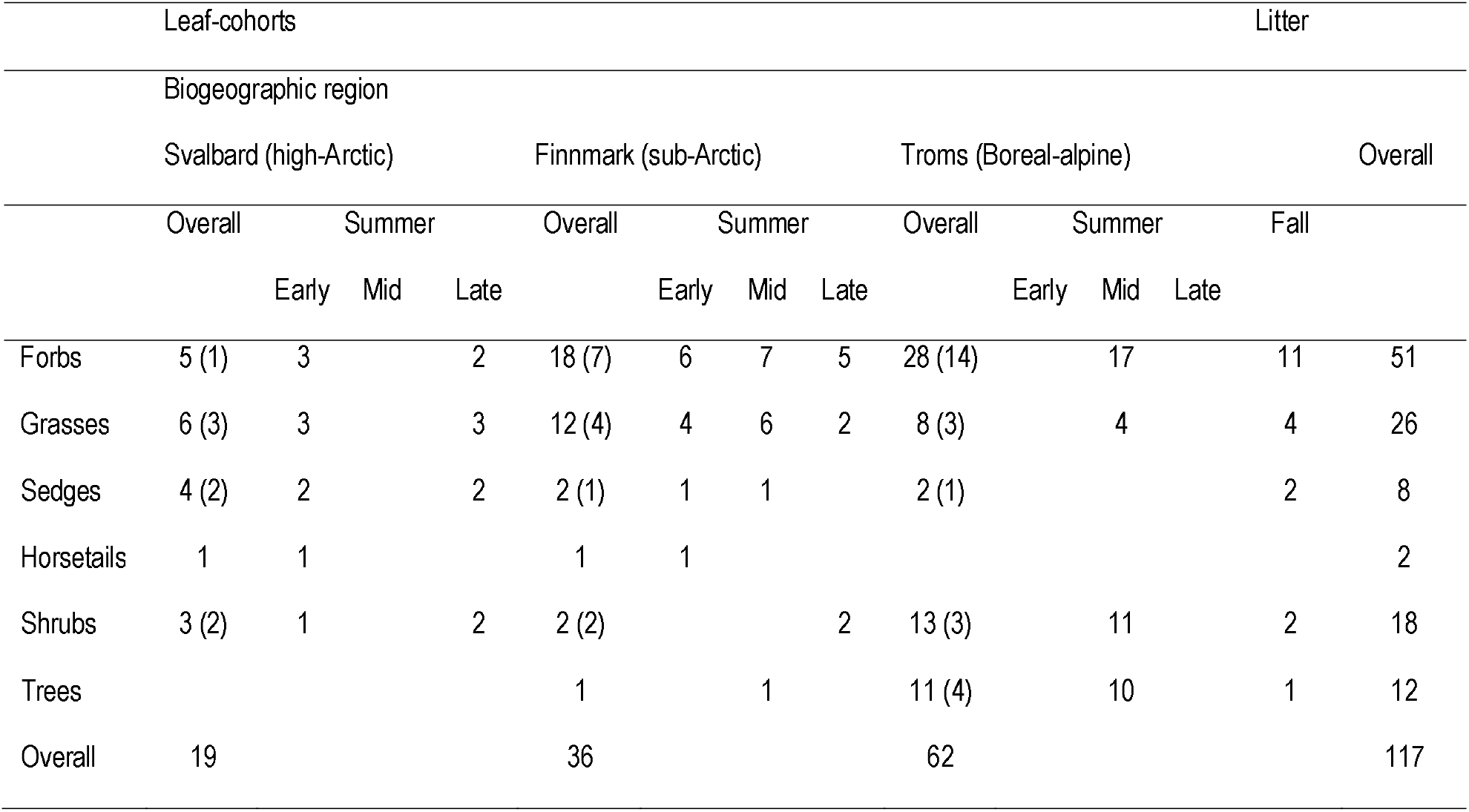
Overview of leaf-cohorts (set of single leaves merged into tablets) sorted according to biogeographic region and growth forms, and further split into phenological stage as indicated by sampling in early, mid and late summer, and including an overview of litter-cohorts all sampled in fall. Numbers in parenthesis refer to number of unique species within each growth form. More information about the cohorts is provided in the data overview (https://opendata.uit.no).

In addition to assessing if the arctic-alpine calibration models can be applied to spectra derived from whole leaves, we also assessed the number of spectra needed for predicting accurate chemical content in whole, single leaves as a guide to future sampling. For this purpose, we sampled fresh leaves from the Varanger Peninsula in Finnmark (70° N, 30° E) during the summer season of 2018. We sampled 22 single leaves of different leaf sizes from a total of 18 species, representing forbs, grasses and shrubs (Table S1), and sorted them in size classes of small leaves (Ø < 1 cm), medium leaves (Ø between 1 and 3 cm) and large leaves (Ø > 3 cm).

### Sample processing

Leaves were sampled individually and immediately put in teabags, pressed dry between filter papers for at least 72 h and dried at 60°C for at least 24 h. In a few cases when we did not have immediate access to plant press and oven-facilities, sampled leaves were stored as dry as possible and pressed at the latest during the evening of the sampling and oven-dried within 5 days.

Per cohort we sampled leaves for a total of approx. 100 mg, which is a leaf mass large enough for making a tablet. The final number of leaves per leaf-cohort was on average 17.29, but varied dependent on both the leaf size of the species and the biogeographic region (Table 2). First, we sampled NIRS-spectra from whole leaves. From the leaf-cohorts spectra were sampled separately from each single leaf, whereas from the litter-cohorts, leaves were stacked and NIRS-spectra were sampled from the leaves collectively. After sampling of spectra from whole leaves, all leaves within a cohort were merged and milled into fine powder using a ball mill (Mixer Mill, MM301; Retsch GmbH & Co. Haan, Germany) and pressed into tablets (Ø 16 mm, 1 mm thick) using a hydraulic press with 4 tons of pressure. Finally, we sampled spectra from each tablet.

**Table 2.** Overview of number of leaves per leaf-cohort and number of spectra sampled per leaf sorted according to biogeographic region.

|  | Biogeographic region |  |  |  |  |  |
| --- | --- | --- | --- | --- | --- | --- |
|  | Svalbard (high-Arctic) |  | Finnmark (sub-Arctic) |  | Troms (Boreal-alpine) |  |
|  | Range | Mean | Range | Mean | Range | Mean |
| Number of leaves per cohort | 20 to 140 | 48 | 1 to 22 | 8.92 | 3 to 126 | 10.57 |
| Number of replicates of spectra per leaf | 3 to 9 | 4.96 | 1 to 12 | 2.47 | 1 to 11 | 1.34 |

Because water shows strong absorption patterns in the near infra-red region (Givens, De Boever & Deaville 1997) both the whole leaves and the tablets were oven-dried for 2 h at 60°C to remove any potential water films, after which they were stored in a desiccator at room temperature (approx. 20°C) until the sampling of spectra.

### Spectral measurements

All spectra were recorded with a portable NIRS spectrometer (FieldSpec 3, Asd Inc., Boulder, Colorado). Spectra of whole leaves were recorded using a custom-made adaptor that can be attached to the ASD Contact probe (Asd Inc., Boulder, Colorado) and allows for measurements of an area as small as Ø 4 mm (Picture 1). The adaptor was made using Delrin, a non-absorptive material similar to that of the original plant adaptor (advice communicated by Asd Inc., Boulder, Colorado). Spectra of tablets were recorded using a similar setup but with an adaptor for an area of Ø 16 mm, exactly matching the size of the tablets (Smis *et al.* 2014).

**Picture 1.**
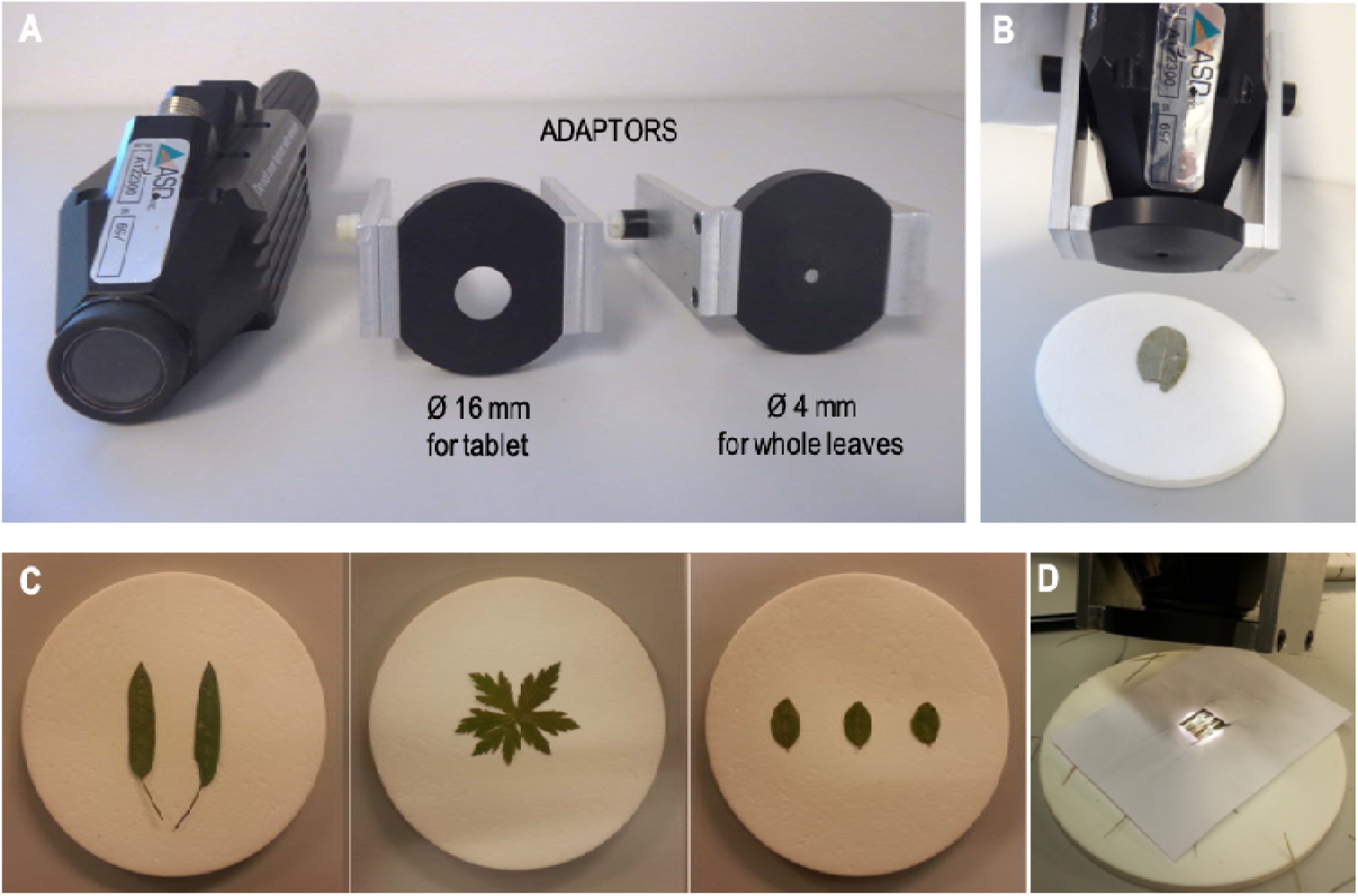
**A.** The plant probe along with custom-made adaptors with a Ø 16 mm area and a Ø 4 mm area for measuring a NIRS-spectrum of a tablet or a single, whole leaf. **B.** The Ø 4 mm adaptor attached to the plant probe ready for scanning a leaf placed on a white pad (a Spectralon) of Ø 10 cm. **C.** Leaves of *Bistorta vivipara, Geranium sylvaticum* and *Vaccinimum myrtillus* prepared for scanning. **D.** Narrow grass leaves prepared for scanning.

Spectra were recorded with monochromatic radiation in the wavelength range of 350-2500nm with NIR, SWIR1 and SWIR2 sensors. The spectra were interpolated to 1 nm intervals based on recordings every 1.4 nm in the 350-1050 nm region and every 2 nm from 1050 to 2500 nm. Wavelength regions for which the different sensors overlap (i.e. 350-380 nm, 760-840 nm, 1700-1800 nm and 2450-2500 nm), were removed from the dataset due to potential inaccuracy in readings. The visible part of the spectrum(380-720 nm) was also removed because this wavelength region has absorption features that might emphasize leaf structural differences (Serbin *et al.* 2014).

The number of sampled spectra per single, whole leaf was dependent on the leaf area, causing a range in spectra sampled (Table 2). Overall, we sampled from 1 to 12 spectra with an average number of 3.53 replicate spectra per leaf and an average number of 61.03 replicate spectra per leaf-cohort, and 14.1 replicate spectra per litter-cohort. For the tablets the average was 3 and 4 replicate spectra for fresh leaves and leaf litter respectively. For the assessment of the number of spectra needed for predicting accurate content of chemical constituents in single leaves, we sampled 10 replicate spectra from small leaves, an average of 14.5 replicate spectra per medium-sized leaves and an average of 32.5 replicate spectra per large-sized leaves. Each spectrum was recorded as absorbance (log 1/ R, where R = reflectance) and the chemical content predicted using the arctic-alpine models for Nitrogen, Phosphorus, Carbon (Murguzur *et al.* 2019) and Silicon content (Smis *et al.* 2014).

### Assessment of method performance

We used the predictions of the chemical content of the tablets (for which the arctic-alpine calibration models are developed) as blueprint to which the predicted chemical content of whole leaves was compared. For the whole, fresh leaves we first calculated the average predicted chemical content per single leaf, after which we calculated the average content per leaf-cohort. For the leaf litter we calculated the average predicted chemical content per litter-cohort directly. We compared the chemical content per cohort as predicted from whole leaves to the chemical content of the cohort as predicted from its tablet. We used linear regression models to assess prediction fit (intercept and slope) and prediction accuracy (root-mean-square error of prediction (RMSEP) and coefficient of determination (R2)). All cohorts were included in the linear regression models for the predictions of Nitrogen, Phosphorus and Carbon content. For the prediction of Silicon content only the silicon-rich growth forms were included (horsetails and graminoids), for which also the arctic-alpine model of Silicon performs best (Smis *et al.* 2014). Negative predictions of Phosphorus content from tablets of two of the cohorts were adjusted to a Phosphorus content of 0.04 % dry weight, the minimum content included in the arctic-alpine calibration model of Phosphorus and measured with chemical analysis (Murguzur *et al.* 2019). The regression analysis was also conducted for fresh leaves and leaf litter separately. The final model was based on all cohorts only if this model was equal or better in accuracy to that of the fresh leaf model, otherwise two separate models (i.e. one for leaf-cohorts and one for litter-cohorts) are presented.

For the assessment of chemical content variation among single leaves within leaf-cohorts, we first corrected predicted values using correction factors achieved from the regression analyses described above. For the chemical constituents where the fit between predicted content from the whole leaves vs the tablets was not 1:1, we applied the intercept and slope as correction factors to adjust the predicted content per leaf. After correction the predicted Phosphorus and Silicon content was negative for a few leaves. These leaves were given a minimum value of content equal to 0.01 % Phosphorus and 0.1 % Silicon.

We assessed intra-cohort variation using a subset of the samples. For intra-cohort variation in chemical content we used leaf-cohorts of Bistorta vivipara, the only species represented with cohorts from all the three biogeographic regions as well as several phenological stages per region. For intra-cohort variation in stoichiometric ratios we used graminoid cohorts sampled from Svalbard in the late season, representing a range of genera for which the predicted chemical content was based on at least four sampled spectra and for which we could include Silicon content. We assessed whether stoichiometric ratios would be more accurately predicted using calibration models based on stoichiometric ratios directly. We made a calibration model for the ratio between Nitrogen and Carbon (Figure S1A). A comparison between the stoichiometric ratios derived from the arctic-alpine calibration models and the new stoichiometric calibration model indicated they were equally precise (Figure S1B), and we proceeded with the arctic-alpine calibration models.

To estimate the number of NIRS-spectra necessary to accurately predict the chemical content of single, whole leaves we sampled a minimum of 10 and a maximum of 42 spectra per leaf. First, we predicted the chemical content from each single spectrum using the arctic-alpine calibration models. Next, we averaged these predictions between an increasing number of replicates (average of predictions from the two first replicate spectra, the three first replicate spectra and so on up to the maximum number of replicate spectra for each leaf). Finally, we compared these averages by calculating their differences. We repeated this procedure 10 times, from each of 10 randomizations of the order in which the spectra were taken. We plotted the differences in predictions as a function of the number of replicate spectra. Based on a graphical presentation of the differences we assessed at what number of replicate spectra the difference in predictions levelled off, with differences approaching zero considered the number of spectra required for accurate predictions of chemical content in single, whole leaves.

All statistical analyses were run in the R environment version 3.4.4 (http://www.r-project.org) using ggplot2 for all graphical presentations.

## Results

The range of chemical content derived from sampled NIRS-spectra of milled and tableted leaves with the Ø 16 mm plant adaptor (Picture 1) was considerable (Table 3A), providing a range in chemical contents for which to pursue the comparison between predictions from whole leaves and tablets.

**Table 3.** Chemical content of milled and tableted leaf- and litter-cohorts as predicted by the arctic-alpine models (Smis *et al.* 2014; Murguzur *et al.* 2019) and applied in this study (A). Model parameters for the regression analysis between chemical content predicted from single leaves and tablets (B). For Phosphorus and Carbon the model parameters improved when separating leaf- and litter-cohorts, whereas for Nitrogen and Silicon the best model included both cohorts. *The Silicon model only includes graminoids and horsetails as these are growth forms with higher Silicon content.

|  |  | A Content (% dry weight) |  | B Model parameters |  |  |  |
| --- | --- | --- | --- | --- | --- | --- | --- |
| Chemical trait |  | Mean | Range | Intercept | Slope | R<sup>2</sup> | RMSE |
| Nitrogen |  | 2.100 | 0.032 - 4.515 | 1.073 | 0.604 | 0.88 | 0.824 |
| Phosphorus | Leaf-cohorts | 0.184 | 0.040 - 0.443 | -0.014 | 0.842 | 0.65 | 0.081 |
|  | Litter-cohorts | 0.127 | 0.040 – 0.291 | -0.003 | 1.165 | 0.56 | 0.053 |
| Carbon | Leaf-cohorts | 45.35 | 40.07 - 51.84 | 7.165 | 0.811 | 0.78 | 2.199 |
|  | Litter-cohorts | 46.48 | 43.07 – 55.19 | -14.403 | 1.341 | 0.91 | 1.654 |
| Silicon* |  | 0.991 | 0 - 2.489 | 0.612 | 0.699 | 0.67 | 0.677 |

We found the arctic-alpine calibration models performed well in predicting content of chemical traits of whole leaves. Predictions of chemical content of a cohort when based on spectra sampled from leaves (using a Ø 4 mm plant adaptor) correlated well with that of predictions based on sampled spectra from the same leaves as milled and tableted (the standardized way of preparing leaf material for measurement of chemical content using NIRS) (Figure 1, Table 3B). For Nitrogen and Silicon, we found both leaf- and litter-cohorts were fitting in a common model (Figure 1). For Phosphorus and Carbon, we found the slope of the litter-cohorts was steeper than that of the leaf-cohorts (Figure 1). Overall, the predicted content from whole leaves differed in range to that of the predicted content from tablets (Figure 1), and the intercept and slope of the regressions deviated from an ideal relationship of 1:1 for all the chemical traits (Table 3). Hence, in order to achieve actual chemical content predictions from the Ø 4 mm sampled spectra of whole leaves, the initial predictions from the arctic-alpine calibration models must be corrected.

**Figure 1.**
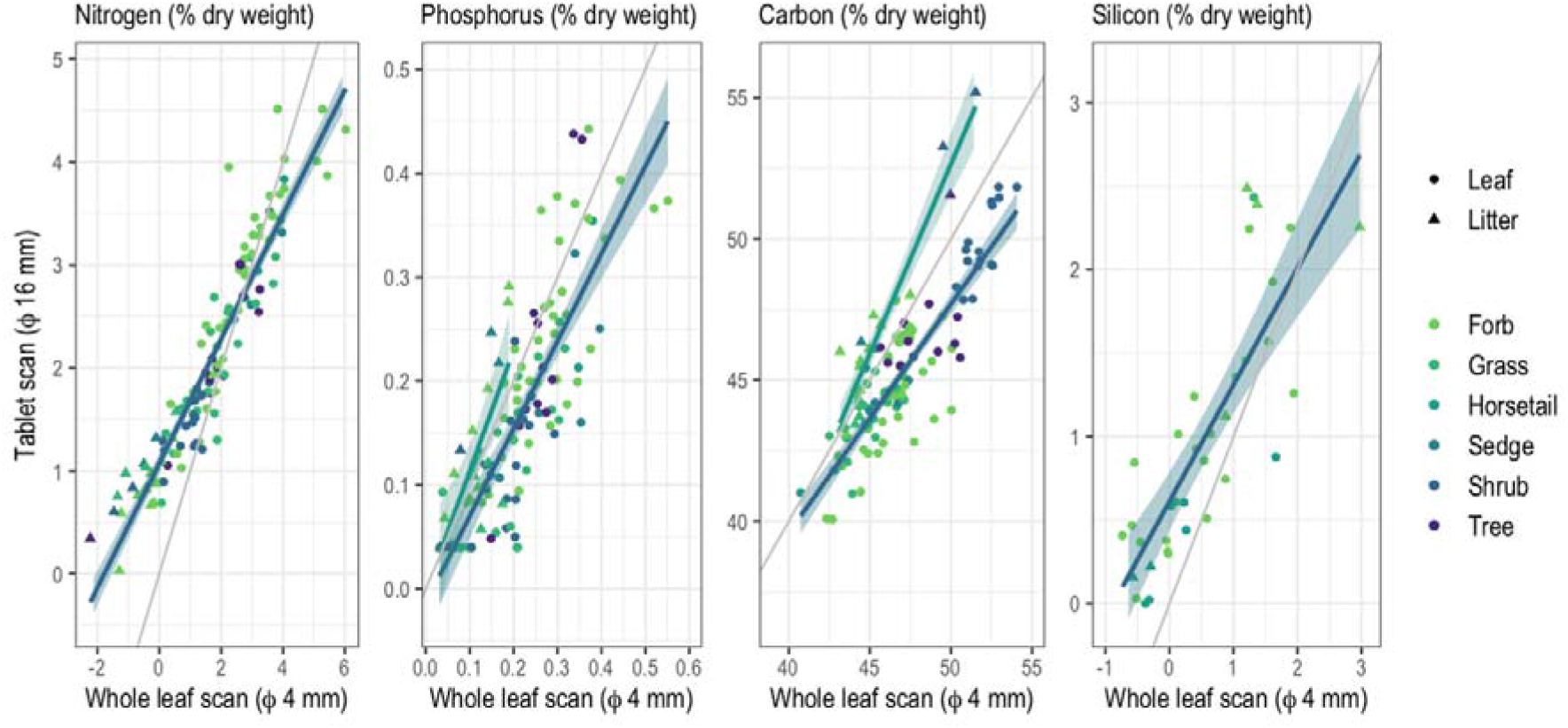
Relationships between Nitrogen, Phosphorus, Carbon and Silicon content of whole leaves and tablets, separate for leaf-cohorts (Leaf) and litter-cohorts (Litter) when their separation improved the linear regression models. The correlation for Silicon content is based on silicon-rich growth forms only. The grey line shows the ideal 1:1 relationship.

The predicted chemical content of single leaves within Bistorta leaf-cohorts showed a considerable variation (Figure 2). The range in chemical content among leaves within a cohort was particularly large for the cohorts from Svalbard (Figure 2, Figure S2), and with a larger range in chemical content in early as opposed to late season. In general, the range in predicted chemical content among leaves within cohorts was equal to or larger than the range in predicted content among seasons and biogeographic regions, as indicated by the predicted content of tablets.

**Figure 2.**
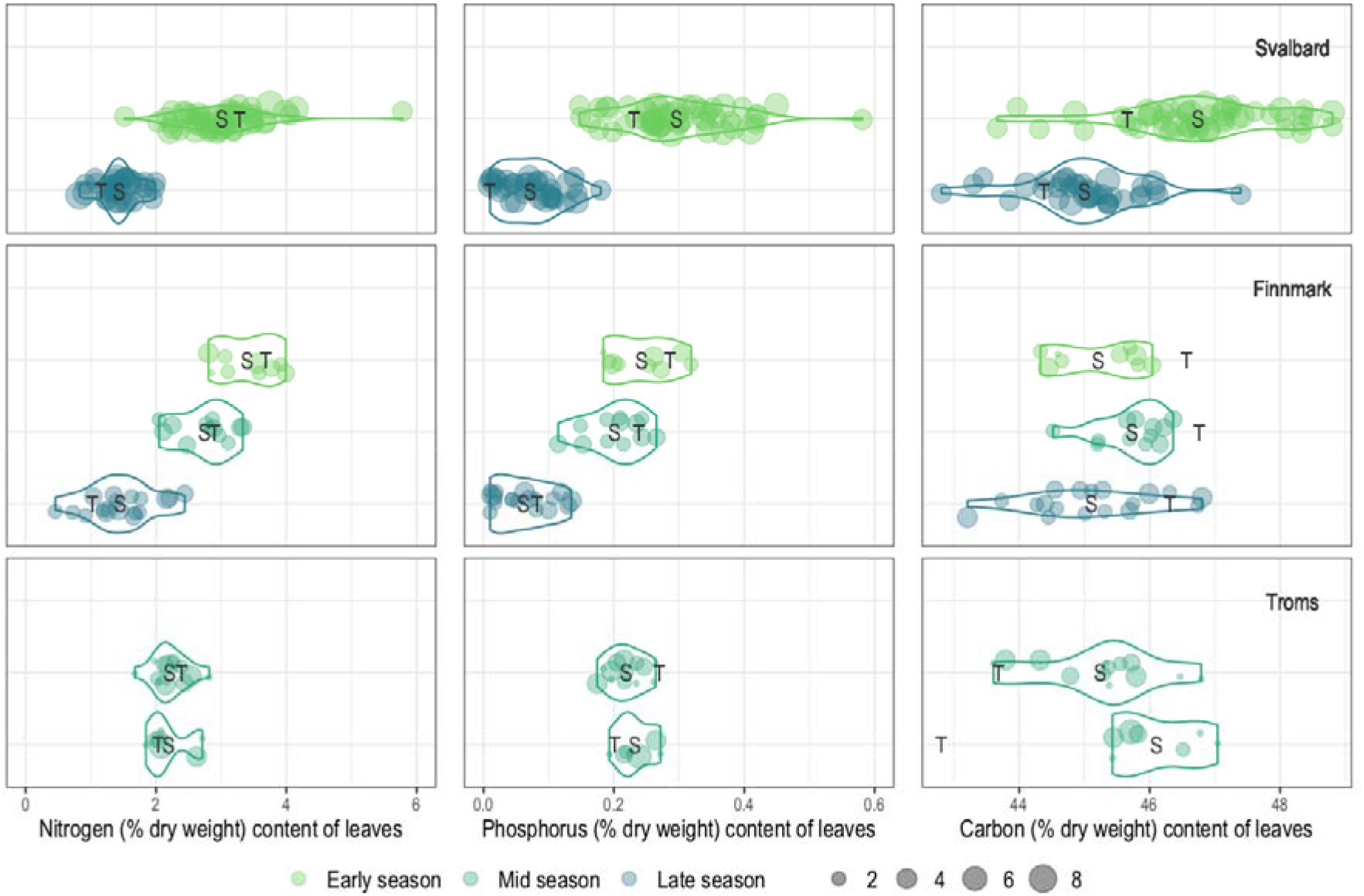
Violin plots of foliar content of Nitrogen, Phosphorus and Carbon (% dry weight) of leaf-cohorts of *Bistorta vivipara*. Each violin represents one leaf-cohort and each dot within each violin represents the chemical content of a unique, single leaf with the size of each dot representing the number of replicate spectra as basis for the predicted chemical content. Cohorts are sorted according to the biogeographic region (Svalbard, Finnmark and Troms) and the season (early, mid and late) they were sampled. The chemical content of the cohort tablet (T) and the cohort average across all leaves (S) are projected onto its respective violin. The leaves sampled from Svalbard were inherently smaller in size than in the two other regions causing the cohorts to have many more leaves(and hence the dense appearance in the plot).

The single, whole leaf predictions were attained on the basis of several spectra sampled per leaf but for a few leaves from the Bistorta leaf-cohorts of the Finnmark and Troms regions that were based on one spectrum only (Figure 2). The predicted content of tablets was both larger, similar and smaller than that of the average of predicted chemical content of the single leaves (Figure 2; forall cohorts see Figure 1). In particular there was a large discrepancy between leaves and tablets for the predicted content of Carbon in the Bistorta leaf-cohorts from the Troms region, the region with most leaves with only one replicate spectrum.

The predicted stoichiometric ratios of single leaves within the graminoid cohorts from Svalbard showed a considerable variation (Figure 3). In general, the three cohorts of grasses showed most variation among leaves, and especially the grass *Calamagrostis*. The sedge *Eriophorum* showed the least variation among leaves. For most cohorts the average of predicted stoichiometric ratios of the leaves overlapped or were close to that of the tablets.

**Figure 3.**
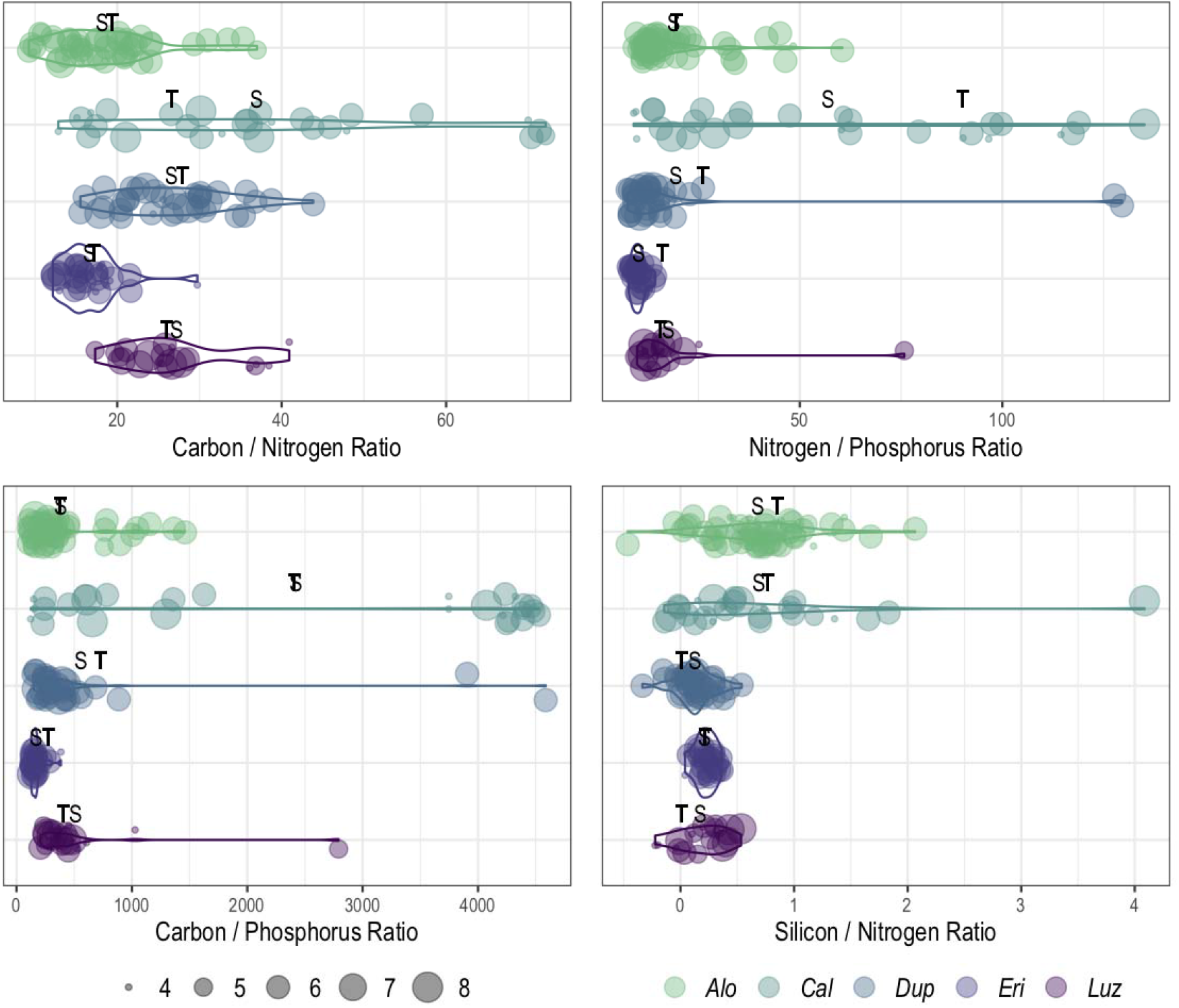
Violin plots of stoichiometric ratios between Nitrogen, Phosphorus, Carbon and Silicon content (% dry weight) of leaf-cohorts of graminoid species sampled from Svalbard in the late season. Each violin represents one leaf-cohort and each dot within each violin represents the stoichiometric ratio of a single leaf, with the size of the dot representing the number of replicate spectra as basis for the predicted stoichiometric ratio. The stoichiometric ratios of the cohort tablets (T) and the average across the cohort leaves (S) are projected onto its respective violin. The graminoid genera included are *Alopecurus (Alo), Calamagrostis (Cal), Duponita (Dup), Eriophorum (Eri) and Luzula (Luz)*.

The precision in predicted chemical content per leaf was dependent on the number of sampled spectra. There was a sharp increase in precision already at 4-5 sampled spectra per leaf, as indicated by a sharp decrease in difference in predictions between 2-3 and 4-5 sampled spectra (Figure 4). When comparing the difference in predicted chemical content to that of the average chemical content of the leaves (insets), the maximum prediction inaccuracy was up to 12.5 % when using only two replicate spectra and dropped to approx. 3 % when using 5 spectra. This supports that a few spectra only can provide an accurate prediction of foliar chemical content of single leaves.

**Figure 4.**
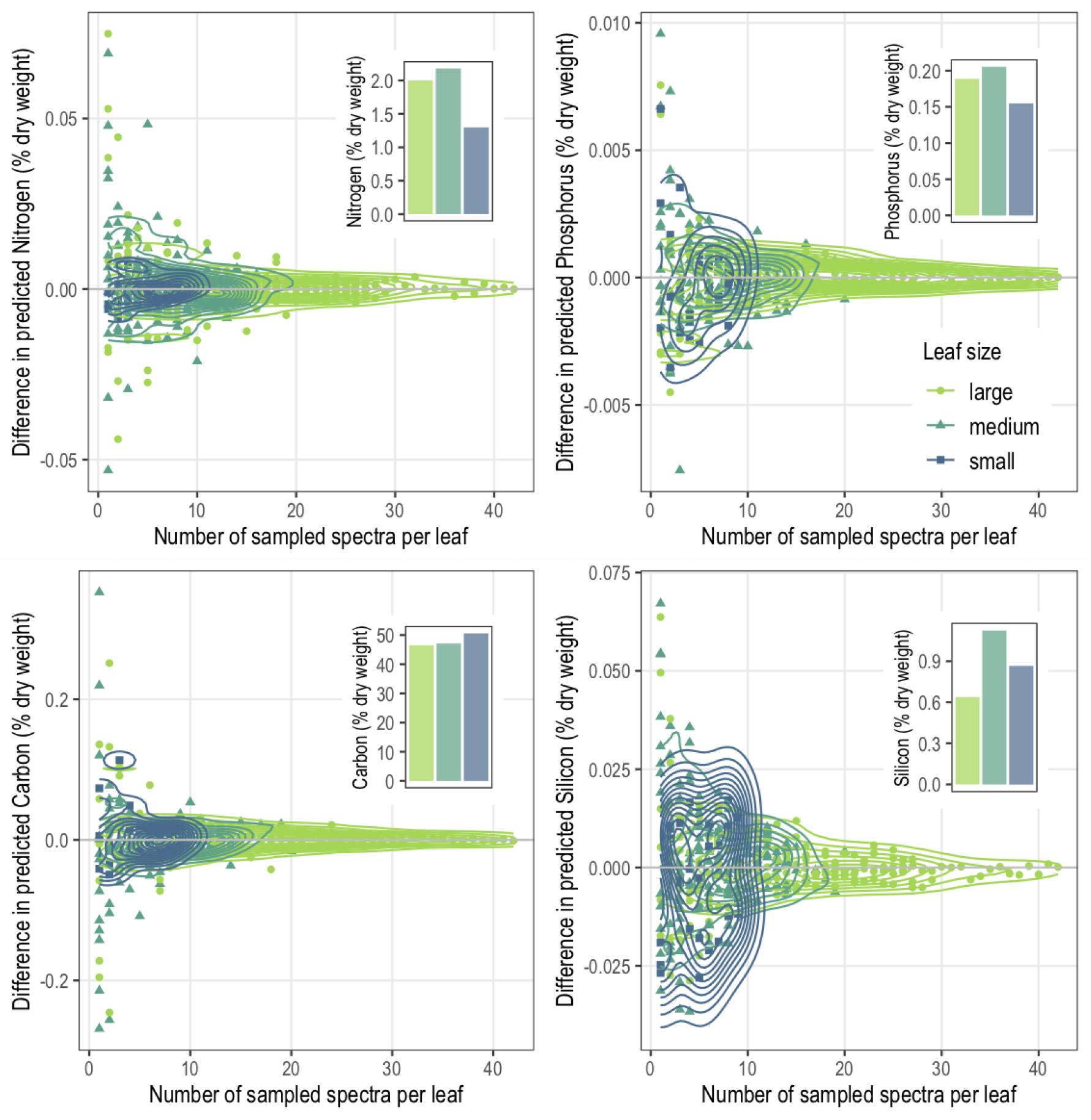
Differences in predicted foliar Nitrogen, Phosphorus, Carbon and Silicon content (% dry weight) as a function of the number of sampled NIRS-spectra per leaf, displayed separately for large-, medium- or small-sized leaves. Each dot represents the average difference in prediction, obtained from randomizing the order of the spectra sampled from a leaf and calculating the differences 10 times (with differences between one and two spectra presented as two sampled spectra, differences between two and three spectra as three sampled spectra a.s.o.). The density-curves demonstrate the overall pattern across all leaves within a leaf-size group. For a comparison to the scale of the chemical content of the leaves, insets show the average chemical content per leaf-size group. Examples of species representing the different leaf-size groups are the grass *Phleum alpinum* and the forbs *Trollius europaeus* and *Soildago virgaurea* for large-sized leaves, the grasses *Anthoxanthum nipponicum* and *Calamagrostis phragmitoides* and the forbs *Bistorta vivipara* and *Rumex acetosa* for medium-sized leaves, and finally the dwarf shrubs *Vaccinium myrtillus* and *Betula nana* for small-sized leaves. In total the leaves of 18 species were included (Table S1).

## Discussion

This study shows that the foliar content of a range of key chemical elements can be measured, using NIRS, from a whole, single leaf and from leaf sizes as small as Ø 4 mm. The NIRS-application is both time- and cost-efficient, and is non-destructive. In a time when marked changes to the environment are happening, and especially in arctic and alpine regions where predicted changes to biogeochemistry are considerable (Jonasson, Chapin & Shaver 2001), we believe our efficient method to achieve chemical traits is a welcomed contribution (Halbritter *et al.* 2019). This is further supported in terms of reduced sampling impact on vegetation when for instance working in experimental or long-term monitoring plots. Also, the quick measure of a range of foliar chemical traits at the level of single leaves opens avenues for research. For instance, chemical traits can be related to that of other traits along with their trade-offs at the level of single leaves, and compared to trade-offs at the level of individuals and populations. Inter- and intra-individual variability in foliar chemical traits can become levels of investigation when studying ecosystem processes such as herbivory and decomposition. In sum, ecological questions for which chemical traits in single leaves are relevant, can easily be addressed through our NIRS-application.

The arctic-alpine calibration models are based on spectral data of milled and pressed plant material. The purpose of milling and tableting leaf material is to create a homogeneous surface and reduce random light scattering (Smis *et al.* 2014). Reduced precision is found when predicting from fresh leaves as opposed to dried and milled leaves, yet the loss in precision does not make predictions from fresh leaves inferior (Couture *et al.* 2016). Furthermore, any gain in information acquired from having time to process more samples (when avoiding tedious processing of leaf samples) may compensate for less accurate predictions (Couture *et al.* 2016). In this study any reduced accuracy in the prediction of chemical trait values from the dried and pressed leaves compared to that of the tablets could not be estimated directly as most single leaves were too small for tableting (and too small for providing wet chemistry measures of the chemical content). However, we found that the accuracy of the measure of chemical content in a single leaf increased with the number of spectra sampled, suggesting the uneven surface of a leaf (such as that of veins and other structures) does not interfere with predictions as long as several spectra are sampled.

The arctic-alpine calibration model of Silicon performs best for Silicon-rich growth forms (Smis *et al.* 2014). Perhaps for this reason we found the model was only applicable to single leaves of Silicon-rich growth forms. Furthermore, species of growth forms with low Silicon content such as forbs, shrubs and trees made up two thirds of all samples (Table 1), hence their inclusion would have caused a bias in the regression analysis towards small content. Also, the predicted Silicon content of single leaves of these Silicon-poor species was sometimes spurious. This indicates aspects of the leaf surface, in turn affecting the spectral signature, interfered with the Silicon-model and that foliar Silicon content of Silicon-poor growth forms are best measured in a homogeneous surface such as that of milled material (Smis *et al.* 2014).

The ability to address chemical traits of single leaves provides the opportunity to assess intraspecific chemical trait variability at several scales, including the within individual variability (Albert *et al.* 2011; Bolnick *et al.* 2011). Indeed, the Nitrogen content related trade-off, or dilemma, of leaves being palatable and efficient in production as opposed to investing in defenses (Díaz *et al.* 2016), may play out differently among single leaves within a plant individual. For instance, plant herbivore interactions between trees and large ungulates can promote changes at both the modular and genetic level (Danell *et al.* 2003). In response to herbivory by moose the deciduous tree Betula pendula allocates more Nitrogen to leaves on shoots browsed by the herbivore than to leaves on lesser-browsed shoots (Danell, Huss-Danell & Bergstrom 1985). There are also several other ways by which intraspecific trait variation - that is variation both within and among individuals of the same species - could alter community structure or dynamics (Bolnick *et al.* 2011). For instance, ecological interactions may depend non-linearly on the variations in a trait, or trait variation may determine the number of ecological interactions taking place (Bolnick *et al.* 2011), hence knowledge of the intraindividual trait variation may increase our predictive ability of ecological interactions. In turn, chemical trait variation among leaves is likely to drive differences in biodiversity among individual plants. The arctic-alpine calibration models, and potentially also other NIRS-based calibration models, provide an opportunity to address such within-individual variation for a range of chemical traits. Furthermore, NIRS-based spectral information at the leaf level also hold the potential for being scaled up to larger scales. Measures at the leaf level within individual plots can be scaled up to canopy, community and landscape levels, and even larger scales, where for each level confounding factors that blur understanding can be addressed. Such scaling also provide efficient measures of biodiversity (Cavender-Bares *et al.* 2017).

There are several aspects of leaves for which a focus on their chemical content may be worth-while. Leaves are functional units for photosynthesis. Leaves are modular units constantly produced and discarded from plant individuals. Leaves are the units often selected for by herbivores. All these functional roles of leaves suggest their chemical content varies, and that measuring their chemical traits at the scale of the functional leaf unit opens avenues to what questions we can ask in ecology. The arctic-alpine calibration models for NIRS-based prediction of foliar Nitrogen, Phosphorus, Carbon (Murguzur *et al.* 2019) and Silicon content (Smis *et al.* 2014) can be applied to achieve chemical traits from single, whole leaves, and as such may be the method to open these avenues.

## Supporting information

Bon et al Supplementary files

## Acknowledgement

We are grateful to Francisco Javier Ancin Murguzur for discussions on NIRS applications and for designing the adaptors, and to Adriaan Smis, Kelsey Lorberau, Mikel Moriano and Jake Robinson for assistance in processing samples. This study received financial support to KAB from the High North Research Centre for Climate and the Environment (The FRAM Centre, www.framsenteret.no) and is an integral study to the Climate-ecological Observatory of the Arctic Tundra, COAT (www.coat.no). There is no conflict of interest to declare for this study.

## Author Contributions Statement

MPB, KAB, TM and SK conceived and developed the study, HB did the modelling, MPB, HB and KAB drafted the paper with help of all authors. All authors gave final approval for publication.

## Data availability

Data will be uploaded to UiT Open Research Data (https://opendata.uit.no) upon acceptance of the manuscript.

