## Supplementary material for "One leaf for all: Chemical traits of single leaves measured at the leaf surface using Near infrared-reflectance spectroscopy (NIRS)": Bon et al Supplementary files

**Table S1.** Overview of samples used to assess the number of spectra needed for predicting accurate chemical content in single leaves (Figure 4). Samples are sorted according to source species and the leaf-size category applied.

| Species | Large leaf  (Ø > 3 cm) | Medium leaf  (Ø >1 U < 3 cm) | Small leaf  (Ø < 1 cm) |
| --- | --- | --- | --- |
| *Anthoxanthum nipponicum* |  | 1 |  |
| *Betula nana* |  | 1 | 1 |
| *Bistorta vivipara* |  | 2 |  |
| *Calamagrostis phragmitoides* |  | 1 |  |
| *Comarum palustre* | 1 |  |  |
| *Chamaepericlymenum suecicum* |  |  | 1 |
| *Deschampsia cespitosa* |  | 1 |  |
| *Geranium sylvaticum* | 1 |  |  |
| *Phleum alpinum* | 1 |  |  |
| *Poa spp* |  | 1 |  |
| *Rumex acetosa* | 1 | 1 |  |
| *Solidago virgaurea* | 1 | 1 |  |
| *Stellaria nemorum* |  | 1 |  |
| *Trientalis europaea* |  | 1 |  |
| *Trollius europaeus* | 1 |  |  |
| *Vaccinium myrtillus* |  | 1 |  |
| *Vaccinium vitis-idea* |  |  | 1 |
| *Viola spp* |  | 1 |  |
| Sum of leaves | 6 | 13 | 3 |

**A**

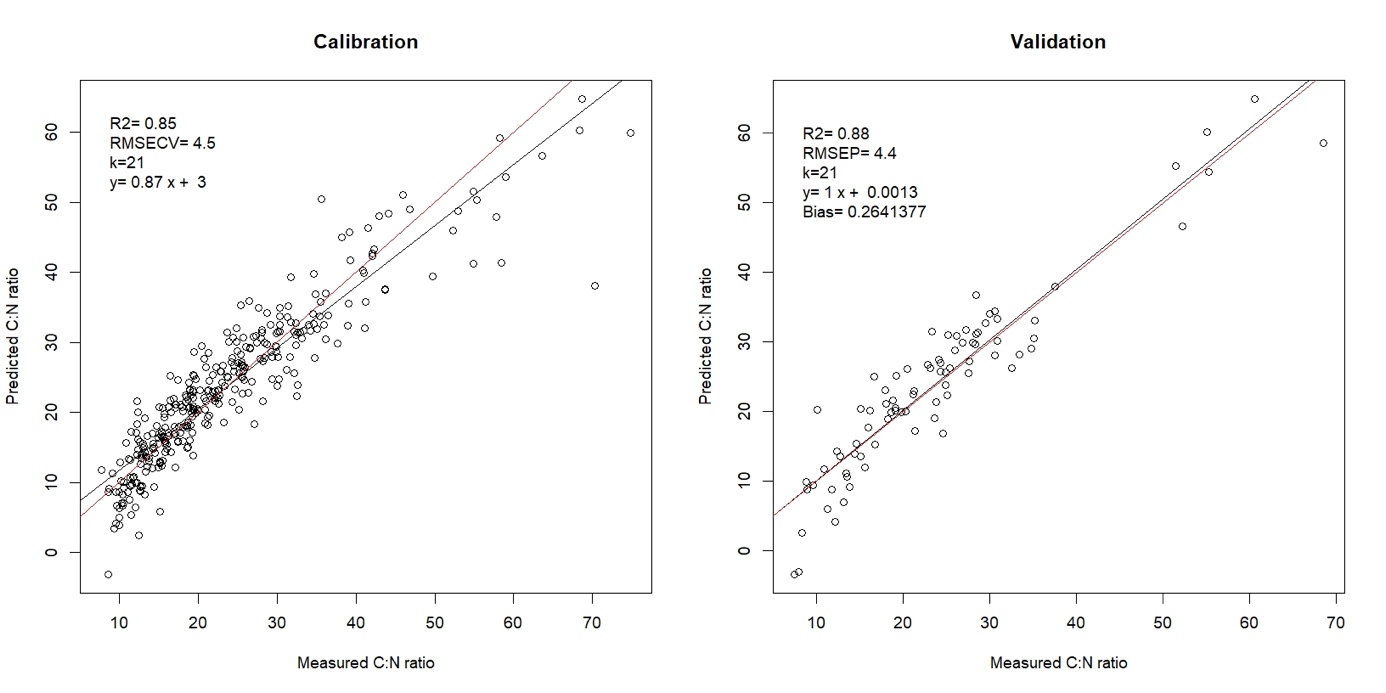

B

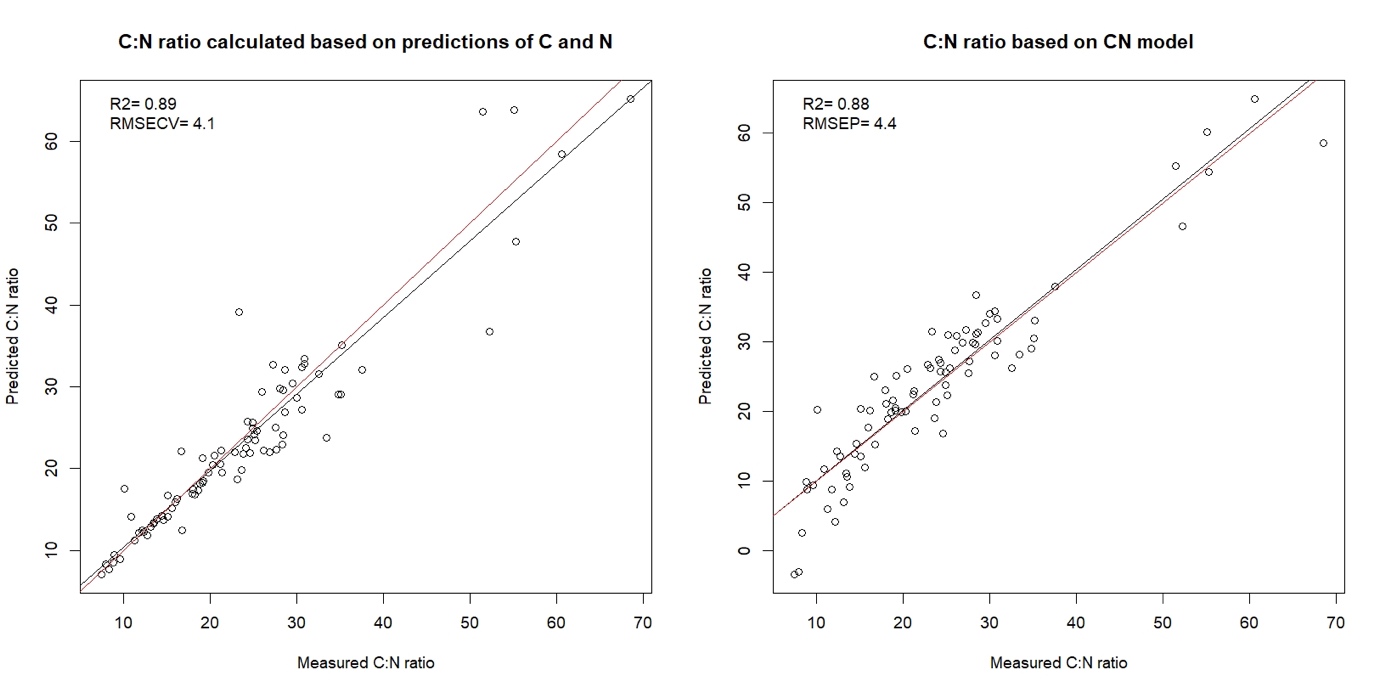

**Figure S1.** Calibration model of the stoichiometric ratio of Carbon to Nitrogen (CN model) and its validation (A). The predicted stoichiometric ratio of Carbon to Nitrogen based on the arctic-alpine calibration models (Murguzur et al. 2019) and the CN model, each compared to the ratio based on chemically measured Carbon and Nitrogen (B).

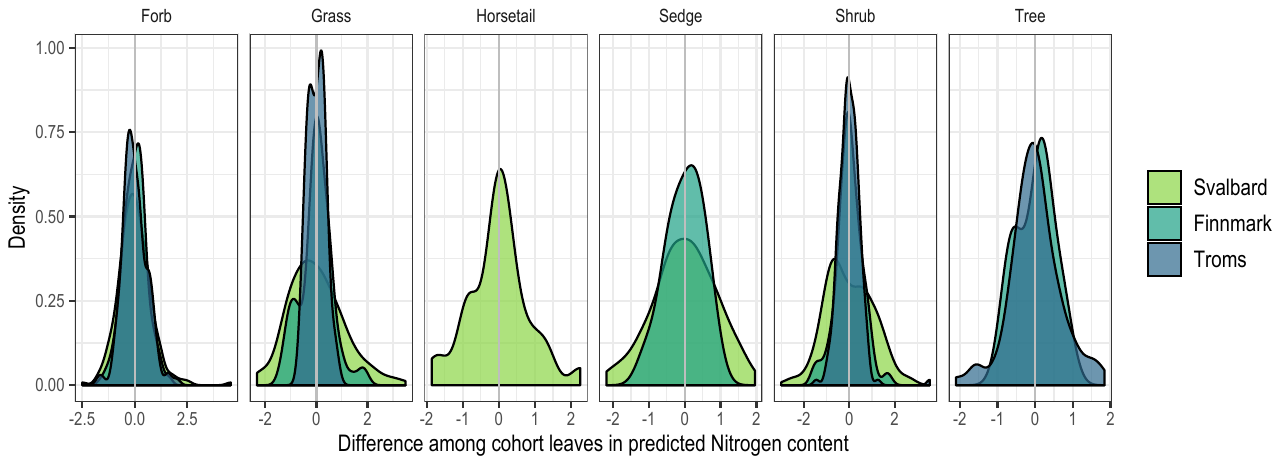

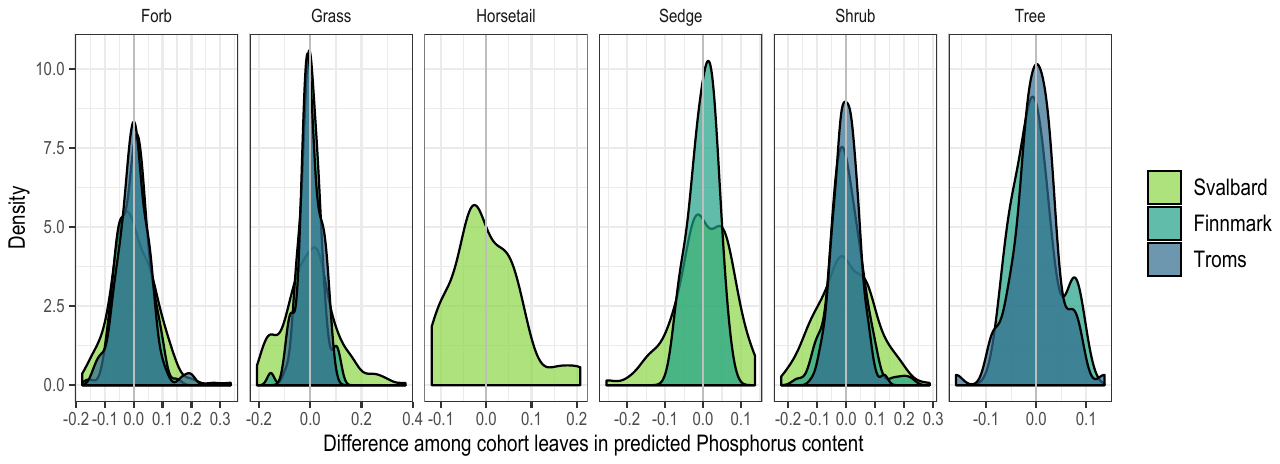

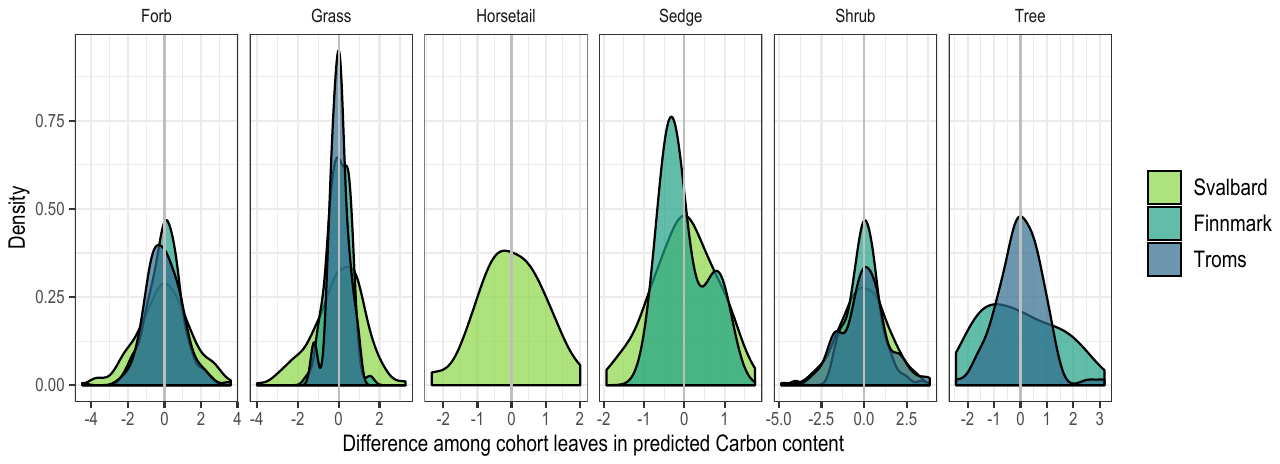

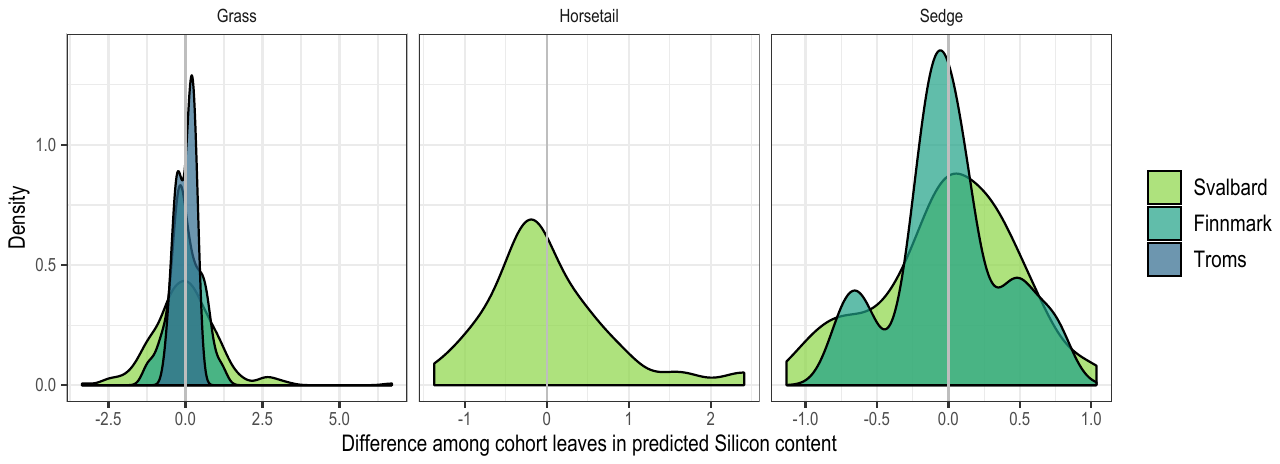

**Figure S2.** Density plots of the standardized predicted chemical content among leaves within cohorts (standardized towards the average content across leaves within the cohorts), sorted according to growth form and biogeographic region.

Murguzur, F.J.A., Bison, M., Smis, A., Böhner, H., Struyf, E., Meire, P. & Bråthen, K.A. (2019) Towards a global arctic-alpine model for Near-infrared reflectance spectroscopy (NIRS) predictions of foliar nitrogen, phosphorus and carbon content. *Scientific Reports,* **9,** 8259.
